# Fluorescent proteins misreport ribosome abundance under translation inhibition

**DOI:** 10.64898/2026.09.17.752513

**Authors:** Aniket Zodage, Justin Chen, Nabigha Mogharbel, Hidde de Jong, Suckjoon Jun

**Author notes:** Corresponding authors : Suckjoon Jun. Contributed equally (Aniket Zodage, Justin Chen).

## Abstract

Fluorescent proteins are widely used as quantitative reporters of protein abundance in living cells. Here we show that this relationship can break down under translation inhibition: the cellular ribosomal abundance can increase while fluorescence intensity remains unchanged or even decreases. Using fluorescent reporters of ribosome abundance in *Escherichia coli* and *Bacillus subtilis*, we find that cellular fluorescence intensity quantitatively tracks ribosome abundance, inferred by RNA-to-protein ratio, under nutrient-limited growth but that is no longer true during chloramphenicol treatment. In *E. coli*, the discrepancy occurs with transcriptional and translational reporters and with both ribosomal protein and ribosomal RNA promoters. To distinguish changes in fluorescence output per reporter from changes in reporter abundance, we constructed a fluorescent protein–LacZ dual reporter in which fluorescence and an independent enzymatic estimate of reporter abundance are obtained from the same protein. Under translation inhibition, LacZ-derived reporter abundance increases whereas fluorescence intensity remains approximately constant, showing that fluorescence output per unit reporter abundance decreases. In a companion study, Bakshi and colleagues demonstrate that the effect extends to other fluorescent proteins and other translation inhibitors, indicating that it is not specific to a particular fluorescent protein or translation inhibitor. Our results show that fluorescent-protein calibration can be condition-dependent and cannot be assumed to transfer across physiological perturbations.

## Introduction

Ribosome abundance has been extensively studied in bacteria because of its central role in growth regulation (*1*–*8*). Early work by Schaechter, Maaløe, and Kjeldgaard (*9*) and by Neidhardt and Magasanik (*2*) showed that, during nutrient-limited growth, ribosome abundance increases approximately linearly with growth rate. Subsequent studies have examined how ribosome abundance and other physiological properties respond to perturbations including transcription and translation inhibition (*3*–*5, 10*), nutrient shifts (*11*–*14*), and temperature shifts (*15, 16*).

Ribosome abundance, commonly defined as the ribosomal protein mass as a fraction of total protein mass, can be estimated from the cellular RNA-to-protein mass ratio. This is because ribosomal RNA (rRNA) constitutes the majority of total cellular RNA, and ribosomal proteins are produced in a stoichiometric proportion to the ribosomal RNA (SI Note 1)(*7, 8, 17*). Mass-spectrometry-based measurements of ribosomal proteins have further supported the use of the RNA-to-protein ratio as a proxy for cellular ribosomal abundance (*5, 10, 18, 19*).

Later, LacZ became a powerful method for more mechanistic studies of *rrn* regulation. For example, Gourse and colleagues placed promoters from one of the seven *rrn* operons, *rrnB*, upstream of *lacZ* at an ectopic chromosomal site and used LacZ activity to characterize *rrnB* promoter regulation under different nutrient conditions in *E. coli* (*20, 21*).

As microscopes have become a convenient and even essential tool in modern biology, fluorescent proteins are a natural candidate to extend LacZ and potentially replace it in most studies. Their advantages are spatiotemporal sensitivity and a wide spectral range, which make them suitable for single-cell studies from protein dynamics to quantitative readout of protein levels (*22*–*24*).

Indeed, several groups including us have recently started to use fluorescent proteins for such purposes: to visualize ribosomes (*25*–*28*), quantify their amount (*29*–*31*), or measure the promoter activities of *rrn*s (*32, 33*). In particular, one of us previously showed that the population-averaged fluorescence intensity of GFP fused to RpsB increased linearly with the nutrient-imposed growth rate, quantitatively reproducing the linear relationship by Scott *et al*. (*3, 31*).

We started this project to expand the use of fluorescent proteins to quantify ribosomal abundance under translation inhibition, a perturbation that led to the discovery of resource-allocation principles in *E. coli* (*3*) and more recently *B. subtilis* (*5*). Despite testing multiple reporter strains, we were unable to reproduce by fluorescence the well-established increase in ribosomal abundance under chloramphenicol treatment. We therefore asked whether the quantitative relationship between fluorescence intensity and reporter abundance itself changes when translation is inhibited.

This question matters because translation inhibition is one of the most basic and widely used physiological perturbations in biology, from the measurement of fluorescent-protein maturation time (*34*) to the study of the cell cycle (*4, 35*) and growth (*3, 5*).

As we show below, one silver lining of our study is that the classic method of Gourse and colleagues, using LacZ under an ectopic *rrn* promoter, appears to be robust under translation inhibition (*20, 21*) even when fluorescent proteins fail the same test. Using an mChartreuse-LacZ fusion, in which both readouts report on the same protein, we further find that the fluorescence output per unit reporter abundance decreases under translation inhibition. An obvious implication is that absolute protein quantification based on fluorescent proteins should be done with caution.

A companion study by Bakshi and colleagues establishes that this effect extends to other fluorescent proteins and other translation inhibitors (*36*) (Refer to Table 5). Together, the two studies show that the loss of quantitative fluorescence under translation inhibition is neither specific to mChartreuse nor to chloramphenicol.

## Results

### Fluorescent reporters accurately report ribosomal abundance under nutrient limited growth conditions

The *E. coli* ribosome is composed of three ribosomal RNA molecules (16S, 23S, 5S rRNAs) and 54 different ribosomal proteins. These molecules assemble into one large ribosomal subunit (50S) and one small subunit (30S) to form a ribosome (70S = 50S + 30S) (Figure 1A).

**Figure 1.**
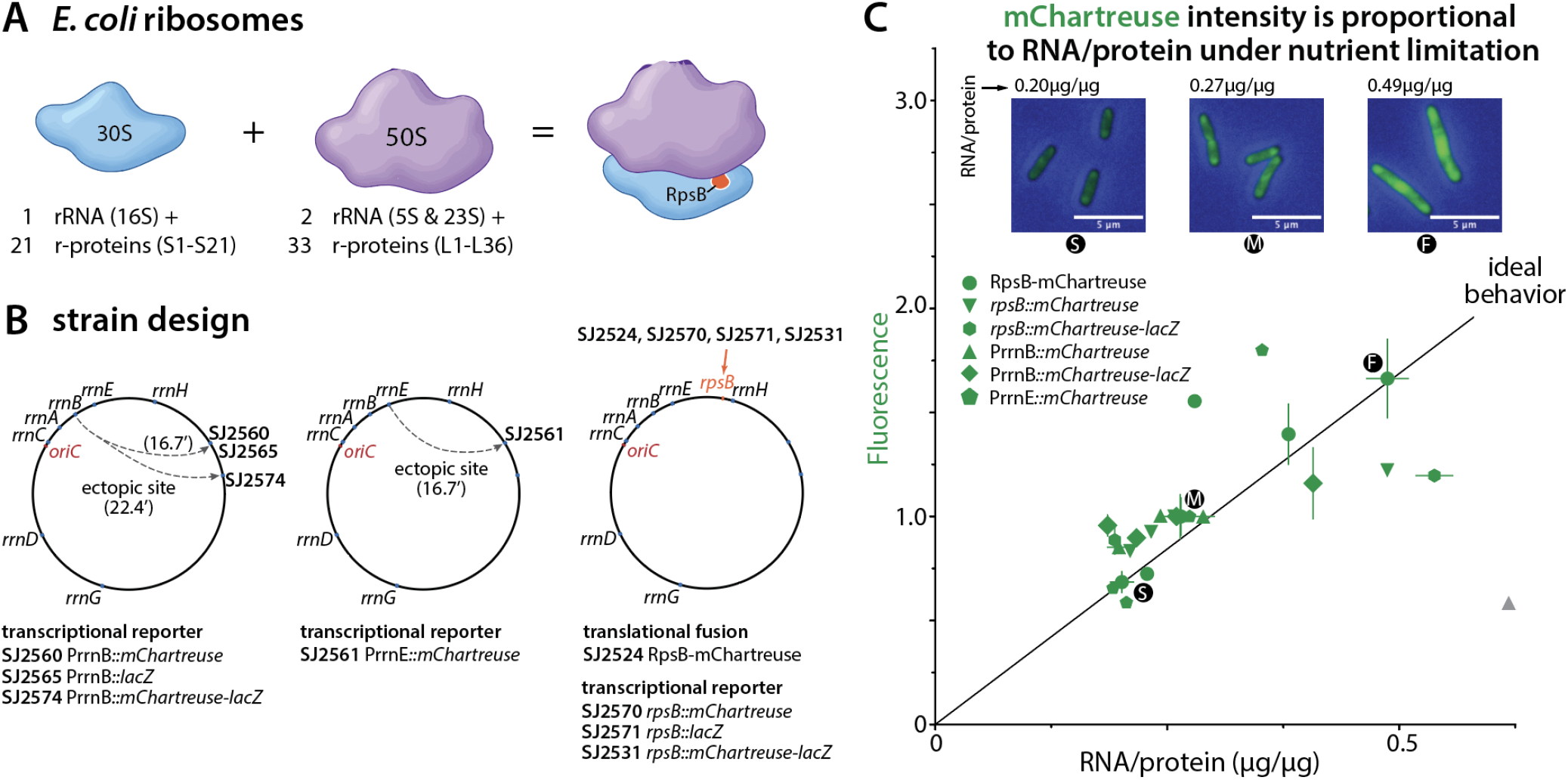
Fluorescent reporter strain designs and their behavior under nutrient-limited growth. (**A**) The *E. coli* 70S ribosome consists of one small (30S) and one large (50S) subunit. The 30S subunit contains 16S Rrna and 21 ribosomal proteins (S1–S21); the 50S subunit contains 23S and 5S rRNAs and 33 ribosomal proteins. RpsB (S2), the fusion partner used in this study, is marked. (**B**) Reporter strain designs and their chromosomal locations. Transcriptional reporters integrated at an ectopic site: PrrnB driving *mChartreuse* (SJ2560), *lacZ* (SJ2565), or *mChartreuse-lacZ* (SJ2574); PrrnE driving *mChartreuse* (SJ2561). Reporters at the native *rpsB* locus: the translational fusion RpsB-mChartreuse (SJ2524), and the transcriptional reporters *rpsB::mChartreuse* (SJ2570), *rpsB::lacZ* (SJ2571), and *rpsB::mChartreuse-lacZ* (SJ2531). Genotypes are listed in Materials and Methods, Table 1. (**C**) Fluorescence intensity (SI Fig 1B) plotted against RNA-to-protein mass ratio (also RNA/protein, SI Fig 1A) measured in the same growth condition. Fluorescence intensity is normalized with the value in MOPS Glucose for each strain, hence it is unitless. The line marked “ideal behavior” is *y* = *a·x*, the direct proportionality expected if fluorescence reports RNA/protein; because fluorescence is unitless, the slope *a* is arbitrary and the line serves as a guide. The agar pad images are for RpsB-mChartreuse strain in MOPS Glycerol (S), MOPS Glucose (M), and MOPS Rich Glycerol (F). The same intensity floor and ceiling were applied across the three images. The grey symbol data is excluded from the linear fit as the fluorescence for that experimental data was significantly lower. Refer to Figure S1B. Growth media are specified in Materials and Methods, Table 2. Error bars are calculated as standard error of mean between at least two replicates. Replicates information is given in Data S1.

For fluorescence measurement of ribosomal abundance, we constructed multiple reporter *E. coli* strains (Figure 1B, Table 1, Materials and Methods) with mChartreuse (*37*) as

**Table 1:** *E. coli* and *B. subtilis* strains used in this study. Sequences used for PrrnB and PrrnE promoters are explained in strain construction below.

| Strain ID | Genotype | Description | Experiments |
| --- | --- | --- | --- |
| *E. coli* |  |  |  |
| SJ81 | MG1655 | MG1655 WT | Used in experiments in Fig. 1 and 2 |
| SJ645 | MG1655 pSIM18 | WT MG1655 strain with pSIM18 lambda-red recombination plasmid | Used for strain construction |
| SJ2524 | *rpsB-mChartreuse kanR* | *mChartreuse* translationally fused to C-terminus of RpsB protein; *kanR* used for selection during lambda-red recombination in SJ645 | Used in experiments in Fig. 1 and 2 |
| SJ2527 | *rpsB::tetA-sacB pSIM18* | Insertion of *tetA-sacB* construct downstream of *rpsB* in SJ645 | Used for strain construction |
| SJ2528 | Δ*lacZYA::kanR* pSIM18 | LacZYA operon deletion in SJ645; pSIM18 lambda-red recombination plasmid is also present | Used for strain construction |
| SJ2529 | Δ*lacZYA::kanR rpsB::tetA-sacB pSIM18* | Insertion of *tetA-sacB* construct downstream of *rpsB* in SJ2528; pSIM18 lambda-red recombination plasmid is also present | Used for strain construction |
| SJ2531 | Δ*lacZYA::kanR rpsB::mChartreuse-lacZ* | Markerless *mChartreuse-lacZ* fusion introduced downstream of *rpsB* in SJ2529 using *tetA-sacB* scarless recombination | Used in Experiments in Fig 1, 2, and 3 |
| SJ2560 | *PrrnB::mChartreuse Pkan::kanR <> tolA* | Cytoplasmic *mChartreuse* reporter driven by the *rrnB* P1-P2 promoter and inserted near *tolA* using in SJ645; *kanR* used for selection | Used in Experiments in Fig 1 and 2 |
| SJ2561 | *PrrnE::mChartreuse Pkan::kanR <> tolA* | Cytoplasmic *mChartreuse* reporter driven by the *rrnE* P1-P2 promoter and inserted near *tolA* in SJ645; *kanR* used for selection. | Used in Experiments in Fig 1 and 2 |
| SJ2565 | Δ*lacZYA::kanR PrrnB::lacZ smR <> tolA* | *lacZ* reporter driven by *rrnB* P1-P2 promoter inserted near *tolA* in SJ2528; *smR* used for selection | Used in experiments in Fig. 1, 2 and 3 |
| SJ2570 | *rpsB::mChartreuse* | Markerless insertion of *mChartreuse* downstream of *rpsB* in SJ2527; using *tetA-sacB* scarless recombination | Used in experiments in Fig. 1 and 2 |
| SJ2571 | Δ*lacZYA::kanR rpsB::lacZ* | Markerless insertion of *lacZ* downstream of *rpsB* in SJ2529 using *tetA-sacB* scarless recombination | Used in experiments in Fig. 1, 2, 3 |
| SJ2574 | Δ*lacZYA::kanR smR PrrnB::mChartreuse-lacZ <> appA* | *mChartreuse-lacZ* reporter driven by *rrnB* P1-P2 inserted within *appA* operon in SJ2528; *smR* used for selection | Used in experiments in Fig. 1, 2, 3 |
| *B. subtilis* |  |  |  |
| JDW94 | YB886 *rpsB-mChartreuse* Δ*hag::MLS* | C-terminal translational fusion of *mChartreuse* with *rpsB*; *hag* gene is deleted to make the strain non-motile | Used in experiments in Figure S2 |

**Table 2:** *E. coli* growth media used in this study.

| Media | Additions | Carbon source and Concentration | Experiments |
| --- | --- | --- | --- |
| MOPS Galactose |  | 0.2% w/v Galactose | Fig 1, 2, 3 |
| MOPS Glycerol |  | 0.4% w/v Glycerol | Fig 1, 2, 3 |
| MOPS Sorbitol |  | 0.2% w/v Sorbitol | Fig 1, 2, 3 |
| MOPS Xylose |  | 0.2% w/v Xylose | Fig 1, 2, 3 |
| MOPS Maltose |  | 0.2% w/v Maltose | Fig 1, 2, 3 |
| MOPS Glucose |  | 0.2% w/v Glucose | Fig 1, 2, 3 |
| MOPS Glucose + 6aa | 6 amino acids (M, H, R, P, T, W) | 0.2% w/v Glucose | Fig 1, 2,3 |
| MOPS Glucose + 12 aa | 12 amino acids (M, H, R, P, T, W, S, L, Y, A, N, D) | 0.2% w/v Glucose | Fig 1, 2,3 |
| MOPS Glucose + CAA | 3% w/v Casamino acids | 0.2% w/v Glucose | Fig 1, 2,3 |
| MOPS Rich Glycerol | EZ rich and ACGU mixture | 0.2% w/v Glycerol | Fig 1, 2, 3 |

**Table 3:** B. subtilis growth media used in this study.

| Media | Carbon source and Concentration | Nitrogen source and Concentration | Experiments |
| --- | --- | --- | --- |
| S7<sub>50</sub> Succinate | 1% w/v Succinate | 0.1% w/v Glutamate | SI Fig. 2 |
| S7<sub>50</sub> Glycerol | 1% w/v Glycerol | 0.1% w/v Glutamate | SI Fig. 2 |
| S7<sub>50</sub> Ribose | 1% w/v Ribose | 0.1% w/v Glutamate | SI Fig. 2 |
| S7<sub>50</sub> Arabinose | 1% w/v Arabinose | 0.1% w/v Glutamate | SI Fig. 2 |
| S7<sub>50</sub> Glucose | 1% w/v Glucose | 0.1% w/v Glutamate | SI Fig. 2 |

- a translational fusion reporter with a small-subunit ribosomal protein RpsB, RpsB-mChartreuse
- a transcriptional reporter of the *rpsB* promoter, PrpsB
- a transcriptional reporter of ectopic ribosomal RNA promoters, *rrnB* P1-P2 promoter (PrrnB), and *rrnE* P1-P2 promoter (PrrnE).

We also constructed LacZ and mChartreuse-LacZ reporters, described below.

Ideally, fluorescence intensity of the reporter protein in all these strains should be proportional to the underlying ribosomal abundance (SI Note 1). This is because the fluorescence intensity is expected to be a proxy of the ribosome concentration. Furthermore, the ribosome concentration is approximately proportional to the ribosomal protein mass fraction, defined here as ribosomal abundance, because the biomass density and protein fraction of cellular biomass remain approximately constant across a range of growth conditions (*38*).

Under nutrient-limited growth, ribosome abundance as inferred from the RNA-to-protein ratio increased linearly with the growth rate for all our fluorescence reporter strains (Figure S1A, Data S1) (*3, 4*). The growth rates were slightly smaller than wild-type MG1655 strains in the previous studies, perhaps due to the additional physiological burden of producing additional proteins using strong promoters. As a result, the slope of RNA-to-protein ratio vs. growth rate was slightly higher (Figure S1A).

The fluorescence intensity for these reporter strains also increased linearly with the growth rate (Figure S1B, Data S1), consistent with Pavlou *et al*. (*31*). Next, we plotted the fluorescence intensity against RNA-to-protein ratio, and found them to be directly proportional to each other (Figure 1C), as previously reported in (*29*). This was seen across all our transcriptional and translational reporter strains.

Similarly, in a *B. subtilis* strain with mChartreuse as a translational-fusion reporter of RpsB, fluorescence intensity increased linearly with the RNA-to-protein ratio (Figure S2) (*5*). These results indicate that both translational-fusion and transcriptional fluorescence reporters track the underlying ribosome abundance under nutrient-limited growth, the relationship being proportional in both *E. coli* and *B. subtilis*.

### Fluorescent reporters exhibit strong deviations under translation inhibition

We next measured mChartreuse fluorescence of our strains under translation inhibition by adding various concentrations of chloramphenicol in the growth medium (Data S1). To our surprise, fluorescence intensity was not proportional to RNA-to-protein ratio (Figure 2A).

**Figure 2.**
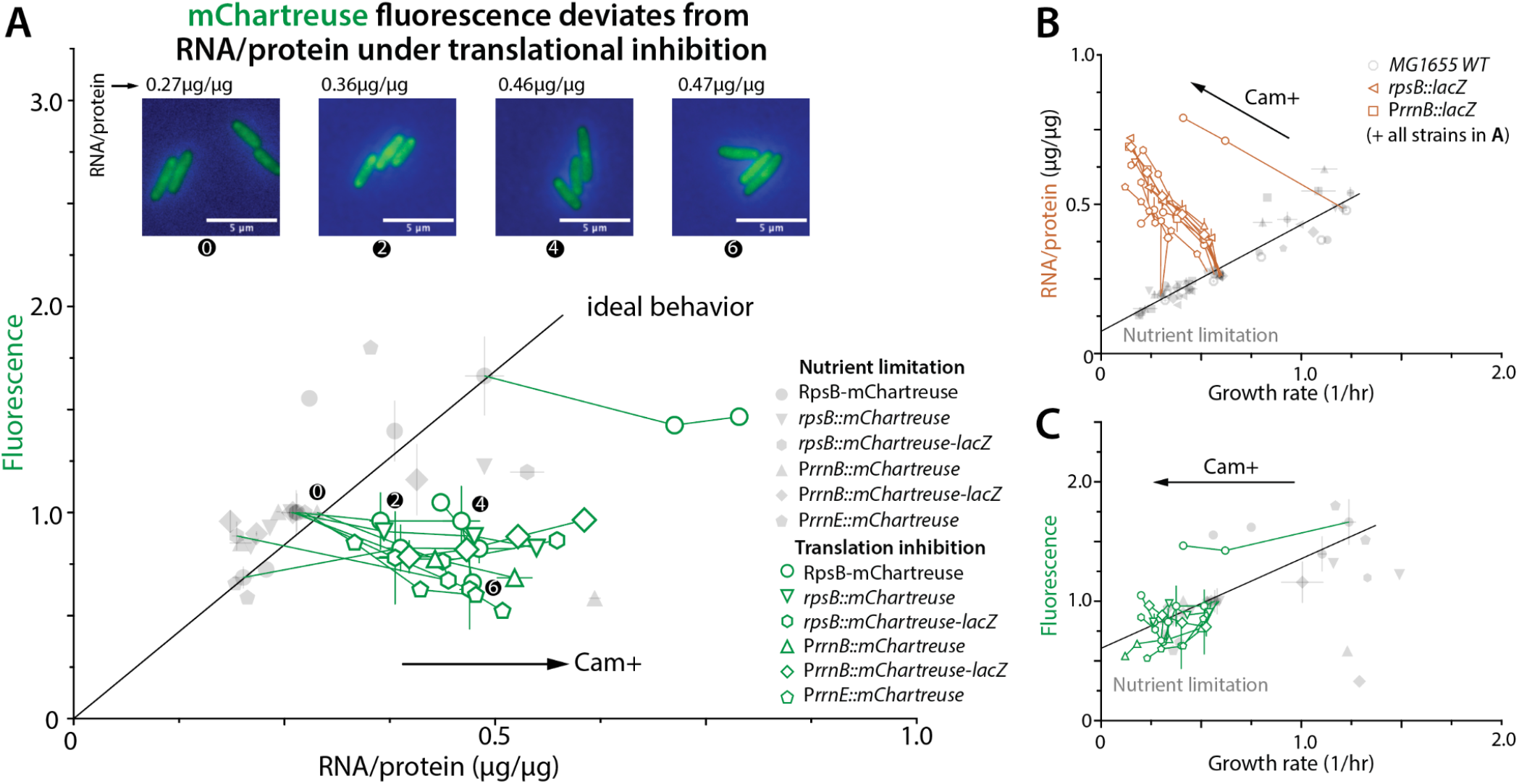
Behavior of fluorescent reporter strains under translation inhibition. Throughout, grey symbols are the nutrient-limitation data of Figure 1, shown for reference, and colored symbols are measurements under translation inhibition. Axis labels are colored to match the data they describe. Connected symbols are a single strain across increasing chloramphenicol concentration, and arrows marked Cam+ give the direction of increasing drug concentration. Chloramphenicol concentrations for each strain are listed in Materials and Methods, Table 4. (**A**) Fluorescence intensity against RNA-to-protein mass ratio (RNA/protein) measured in the same growth condition, for all six mChartreuse reporter strains. Fluorescence intensity is normalized to the value in MOPS glucose for each strain, hence it is unitless. The line marked “ideal behavior” is the direct proportionality expected if fluorescence reports RNA/protein, like in Fig. 1. The agar pad images are for RpsB-mChartreuse strain in MOPS Glucose with addition of 0μM, 2μM, 4μM and 6μM chloramphenicol. The same intensity floor and ceiling were applied for the three images. (**B**) RNA/protein against growth rate, for wild-type MG1655, the *lacZ* reporter strains, and all strains in (**A**). Black line, fit to the nutrient-limitation data. (**C**) Fluorescence intensity (normalized by MOPS Glucose value for each strain) against growth rate, for all strains in (**A**). Black line, fit to the nutrient-limitation data. Error bars are calculated as standard error of mean between at least two replicates. Replicates information is given in Data S1.

This deviation from the expected behavior was due to the decrease in fluorescence intensity under translation inhibition. Specifically, RNA-to-protein ratio increases as chloramphenicol concentrations increase in *E. coli* (Figure 2B) (*3, 4*). However, the mChartreuse fluorescence intensity instead remained approximately constant, or even decreased (Figure 2C). This behavior was seen across all our reporter strains, indicating that this phenomenon is independent of the type of fusion (translational vs. transcriptional) and the promoter identity (ribosomal protein vs. ribosomal RNA promoter).

**Table 4:**
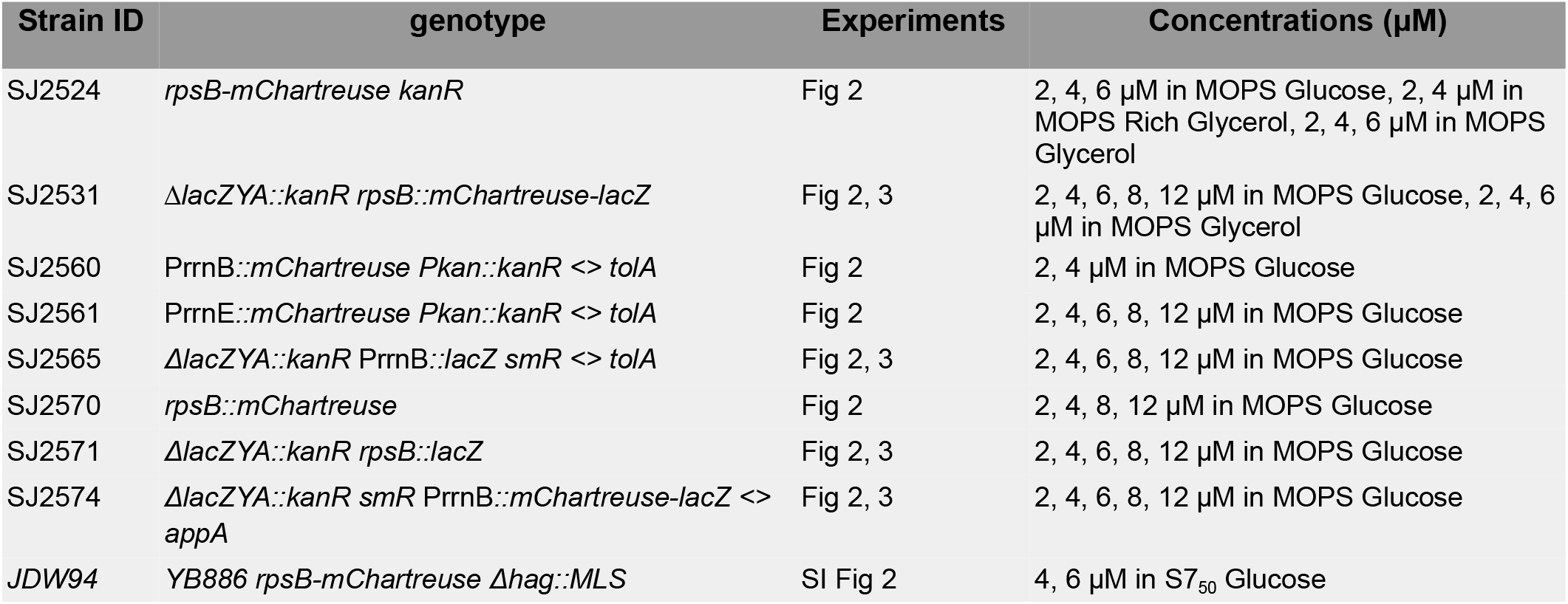
Chloramphenicol concentrations used in this study.

**Table 5.**
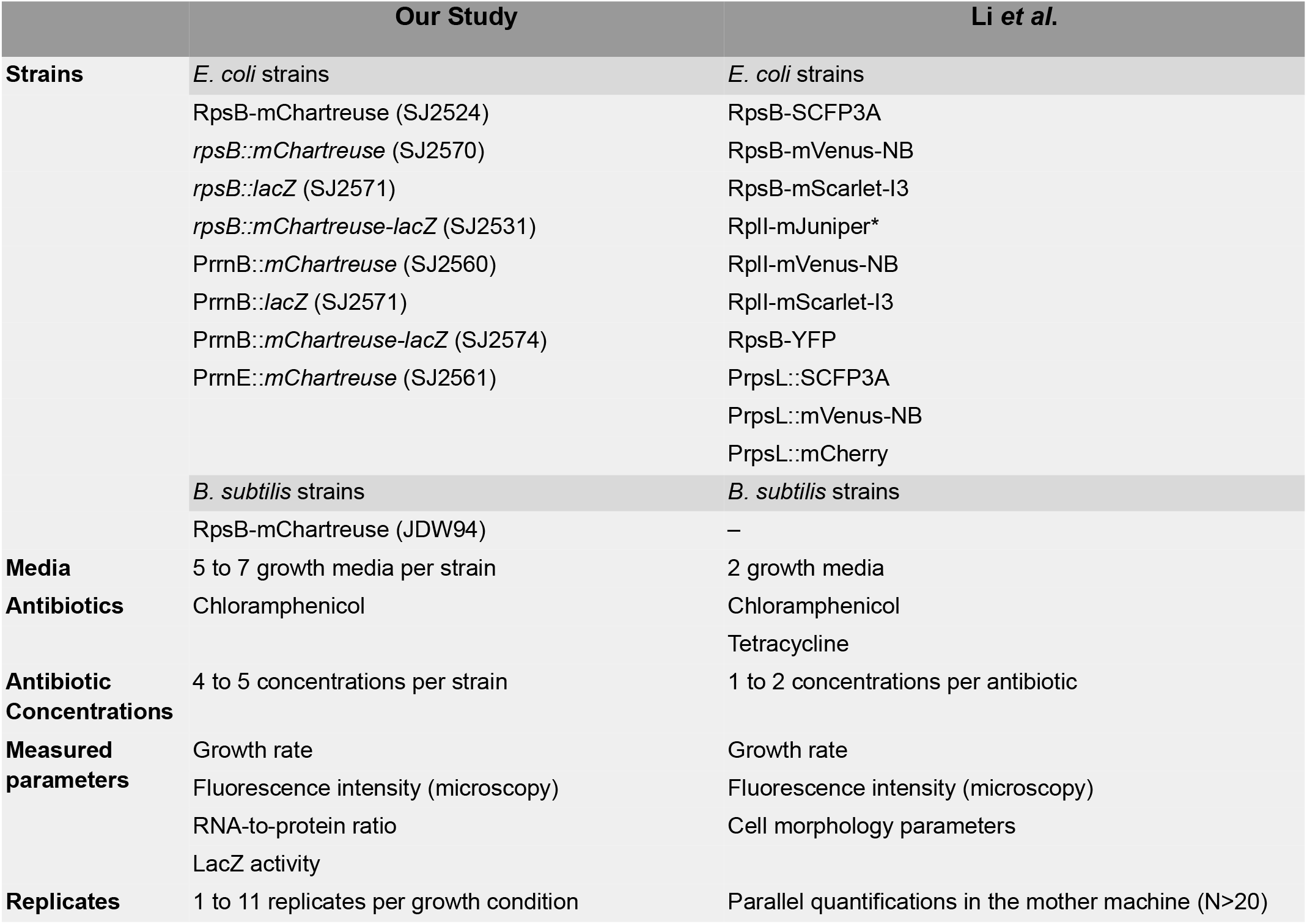
Summary of experiments in the present study and Li *et al*.

*B. subtilis* showed a similar deviation (Figure S2). Under chloramphenicol treatment, RNA-to-protein ratio remains approximately constant in our RpsB-mChartreuse translational fusion strain, as previously reported (*5*). However, mChartreuse fluorescence intensity decreased with increasing chloramphenicol concentration.

Therefore, in stark contrast to nutrient-limited growth, fluorescence reporters misreport ribosome abundance under translation inhibition in both *E. coli* and *B. subtilis*. This discrepancy may arise from a change in fluorescence output of the reporter or from a change in reporter expression. However, from fluorescence alone, we cannot distinguish these two changes and both may contribute. The dual reporter below addresses this directly.

### LacZ under an ectopic *rrn* promoter correctly reports ribosomal abundance

As mentioned earlier, LacZ has been used as a reporter to study ribosomal RNA promoter activity under nutrient limitation (*20, 21*). We thus tested whether LacZ can report ribosome abundance correctly under translation inhibition by constructing two types of strains: PrrnB::*lacZ* and PrrnB::*mChartreuse-lacZ* (Figs. 1B, 3A).

In both strains, we quantified beta-galactosidase activity, which is proportional to the LacZ concentration, from which we calculated LacZ mass as a fraction of total protein mass ([LacZ]) (see Materials and Methods, Data S1). Ideally, by construction, [LacZ] is expected to be proportional to the ribosome abundance (SI note 1). Indeed, we found that [LacZ] is proportional to the ribosome abundance under both growth conditions (Figure 3A), showing that LacZ activities are a quantitative reporter of ribosome abundance, perhaps except under strong translation inhibition with the chloramphenicol concentration close to MIC.

**Figure 3.**
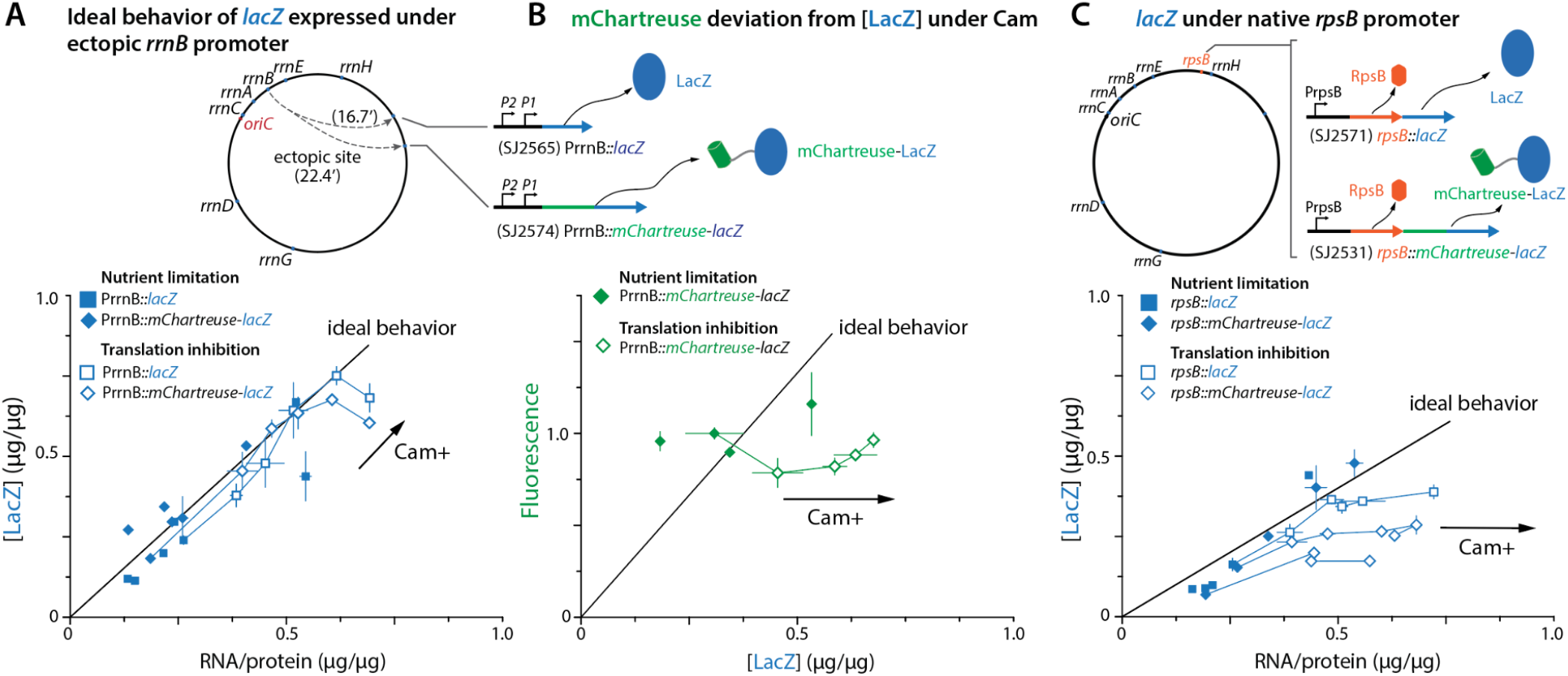
LacZ reporters separate reporter abundance from fluorescence output. (**A**) Top, ectopic PrrnB reporters: *lacZ* alone (SJ2565) and the *mChartreuse-lacZ* translational fusion (SJ2574), under the *rrnB* P1-P2 promoters at the ectopic site. Bottom, [LacZ], calculated as the LacZ mass fraction of total protein mass, against RNA-to-protein mass ratio measured in the same growth condition. Filled symbols, nutrient limitation; open symbols, translation inhibition; the arrow marked Cam+ gives the direction of increasing chloramphenicol. The line marked “ideal behavior” shows direct proportionality between the two quantities. (**B**) Fluorescence intensity against [LacZ] measured from the same mChartreuse-LacZ fusion protein in SJ2574, in the same growth conditions. Because both readouts come from the same protein, their ratio is the fluorescence output per unit reporter abundance. Line as in (**A**). (**C**) Top, reporters in the native *rpsB* operon: *rpsB::lacZ* (SJ2571) and *rpsB::mChartreuse-lacZ* (SJ2531), transcribed from PrpsB downstream of *rpsB*. Bottom, [LacZ] against RNA/protein, plotted as in (**A**). Growth media are specified in Materials and Methods, Table 2. Error bars are calculated as standard error of mean between at least two replicates. Replicates information is given in Data S1.

### Fluorescence output decreases relative to reporter abundance under translation inhibition

Importantly, the mChartreuse fluorescence intensity was not proportional to [LacZ] in the mChartreuse-LacZ translational-fusion dual-reporter strain (Figure 3B, Data S1). Because both readouts come from the same protein, this comparison allows us to compare fluorescence output with an independent estimate of reporter abundance. Under nutrient-limited growth, fluorescence intensity increased proportionally with [LacZ]. Under translation inhibition, fluorescence intensity remained approximately constant while [LacZ] increased. The fluorescence output per unit reporter abundance therefore decreases under translation inhibition, whereas this ratio should be constant for a valid reporter.

mChartreuse and LacZ are encoded on a single transcript and translated as one polypeptide, so effects of translation inhibition on transcription, translation, or reporter expression cannot by themselves explain the divergence of the two readouts. Defects such as premature transcription termination, ribosome stalling, or translation abortion would either reduce both readouts or generate N-terminal mChartreuse-containing products lacking functional LacZ. Such effects would therefore leave fluorescence per unit LacZ activity unchanged or increase it, rather than produce the decrease observed here.

Converting LacZ activity to [LacZ] assumes a specific activity for LacZ (Materials and Methods), but the inference does not require that this specific activity be unchanged. If translation inhibition impairs LacZ folding or activity, the activity-based measurement underestimates how much LacZ is present, and the observed drop in fluorescence per reporter is then conservative. The inference requires only that LacZ specific activity does not increase enough to produce the effect, and the agreement between [LacZ] and RNA-to-protein ratio across the different chloramphenicol concentrations argues against that.

Because fluorescence intensity and LacZ activity are measured from the same expressed protein, changes in the expression of that protein are shared by both readouts and cannot explain their divergence under translation inhibition. Taken together, these results show that the discrepancy arises from the fluorescent output of a reporter rather than from reporter abundance itself.

### LacZ is not a reliable reporter of ribosomal proteins at native promoter

Finally, since LacZ behaved reliably under the ectopic *rrn* promoters either alone or as a translational fusion to a fluorescent protein, we asked whether we can also express *lacZ* under a native promoter of ribosomal proteins. To answer this, we constructed two additional strains and used LacZ and mChartreuse-LacZ as transcriptional reporters of RpsB (Figure 3C).

Surprisingly, and unfortunately, [LacZ] tracked RNA-to-protein ratio under nutrient-limited growth but not under translation inhibition (Data S1). This could be due to the premature transcription termination observed under translation inhibition (*19*), which is consistent with the reduced transcription capacity reported under the same conditions (*33*). Because the position of *lacZ* within the operon for native-site expression is further downstream of the promoter than in the ectopic-site expression, native-site expression would be more affected by premature transcription termination than in the PrrnB reporter strains. However, we did not test this further in this study.

These results show that the robustness of LacZ under the ectopic PrrnB promoter is not a general property of LacZ. Reporter performance depends on the regulatory and genomic context of expression as much as on the reporter molecule itself.

## Discussion

In this report, we examined the use of fluorescent proteins to estimate ribosome abundance. We found fluorescence reporters to be reliable under nutrient limitation, but not under translation inhibition. This was the case regardless of the type of fusion (transcriptional or translational) or the identity of the ribosomal RNA or ribosomal protein promoter. The results of our mChartreuse-LacZ dual reporter strain show that fluorescence output per unit reporter abundance changes under translation inhibition, so fluorescence no longer quantitatively reports how much reporter is present.

A protein reporter differs from RNA-to-protein ratio in one important respect: its expression depends on the very process being inhibited. Translation inhibition therefore perturbs the measuring system as well as the cell, and the extent of this perturbation can depend on the promoter and genomic context of the reporter. Our native-site LacZ constructs illustrate this context dependence, although its origin remains unclear. The dual reporter distinguishes these expression-level effects from the fluorescence-specific discrepancy: because mChartreuse and LacZ are encoded and translated together, defects in reporter expression cannot produce the observed decrease in fluorescence relative to LacZ activity, as shown above.

We do not yet understand why fluorescence output of a reporter decreases relative to its abundance under translation inhibition. This phenomenon is distinct from previously reported problems with fluorescent-protein fusions. An N-terminal YFP tag can generate artefactual MreB helices (*39*), and GFP-FtsZ at elevated expression can inhibit cell division and form abnormal spiral structures (*40*). Fluorescent proteins can also generate apparent intracellular localization through aggregation (*41*). In our case, the dual reporter instead reveals a condition-dependent change in fluorescence output relative to reporter abundance. The molecular basis of this effect remains to be determined.

The case of LacZ is intriguing and important. The ideal behavior was observed only when lacZ was expressed at the ectopic site originally and carefully studied by Gourse and colleagues (*20, 21*). Perhaps these results reinforce the delicate nature of regulation of *rrn* and production of ribosomal proteins, and ultimately ribosome assembly, shaped by evolution.

The mChartreuse-LacZ fusion also offers a general way to validate quantitative fluorescence measurements. Because the fluorescent and enzymatic readouts come from the same expressed protein, an orthogonal reporter of this kind can distinguish changes in reporter abundance from changes in fluorescence output across physiological perturbations.

Our result may also be relevant to measurements of fluorescent-protein maturation, one of the most valuable applications of translation inhibition. Balleza et al. established the systematic framework for measuring maturation times in living cells, using chloramphenicol to arrest synthesis of new fluorescent protein and the subsequent increase in fluorescence to infer the fraction of immature protein and its maturation kinetics (*34*). A constant multiplicative change in fluorescence output that applies equally to the entire fluorescence trace would not affect the normalized maturation kinetics. By contrast, a change in fluorescence output that develops after chloramphenicol addition, on the same timescale as maturation, could distort the inferred maturation curve. Our measurements are at steady state and do not resolve this short-time behavior, so whether this affects maturation-time measurements is worth checking directly on the timescale of the assay.

Fluorescent proteins are extraordinarily useful for studying biological processes in living cells. Our results show that the calibration is a property of the reporter and the condition together, not of the reporter alone. Quantitative fluorescence measurements should therefore be validated under the physiological perturbation in which they are used.

## Materials and Method

### Strains

All the *E. coli* strains used in this study were in the MG1655 background and *B. subtilis* strains in the YB886 background.

### Strain construction

#### E. coli

*E. coli* strains were constructed in the MG1655 background using λ-Red recombination, as described previously (*42*). Recombination was performed using strains carrying the temperature-inducible λ-Red plasmid pSIM18. Linear double-stranded DNA fragments containing the desired fluorescent-protein or *lacZ* reporter constructs, selectable markers where applicable, and approximately 50bp homology arms corresponding to the target chromosomal locus were generated by PCR.

Expression of the λ-Red recombination proteins was induced by incubating cells at 42°C for 15min, followed by cooling on ice for 12min. Approximately 300ng of the appropriate linear DNA fragment was introduced into the induced cells by electroporation. Cells were recovered in LB medium at 37°C for 4hr and plated on LB agar containing the appropriate antibiotic for selection. Candidate colonies were screened by colony PCR, and the intended chromosomal modifications were confirmed by Nanopore sequencing (Plasmidsaurus Inc.).

For markerless reporter fusions, including the *mChartreuse-lacZ* and related constructs, *tetA-sacB* counterselection was used as described previously (*43*). Briefly, a *tetA-sacB* cassette was first inserted at the desired chromosomal locus using λ-Red recombination, and recombinant colonies were selected on tetracycline. The cassette was subsequently replaced with the desired reporter construct through a second round of λ-Red recombination. Cells were plated on LB agar containing 10% sucrose to select against retention of the *sacB* cassette. When substantial background growth occurred, candidate colonies were restreaked on 10% sucrose plates. Final constructs were verified by colony PCR and Nanopore sequencing.

For the *rrnB* P1-P2 promoter, the DNA sequence from -152 to +1 was chosen which includes P1 and P2 promoters. This sequence was previously used by Murray *et al*. (*20*). For the *rrnE* P1-P2 promoter sequence, 554 bp upstream of 16S rRNA in *rrnE* were chosen. This sequence was previously used by Maeda *et al*. (*32*).

#### B. subtilis

The *B. subtilis rpsB-mChartreuse* was constructed using the loop-in/loop-out markerless method described here (*44*). A suicide-vector with *rpsB-mChartreuse* allele, flanked by 500bp homology to the 3’ end of *rpsB* gene, was constructed. This plasmid also has an I-SceI restriction site. This plasmid was transformed into a *B. subtilis* YB886 strain with *hag* deletion. The plasmid integration was selected for using chloramphenicol. Then cells were transformed with a plasmid (with *kanR* selection marker) that expresses the SceI gene constitutively. Then cells were plated on LB+kanamycin. Then resultant colonies were either *B. subtilis* YB886 with *hag* deletion or the *rpsB-mChartreuse* mutants. The colonies were screened using colony PCR and confirmed with sequencing.

### Growth media

We used LB lennox for seed cultures of all the strains (explained in section Culture procedure). Composition of LB lennox is given below

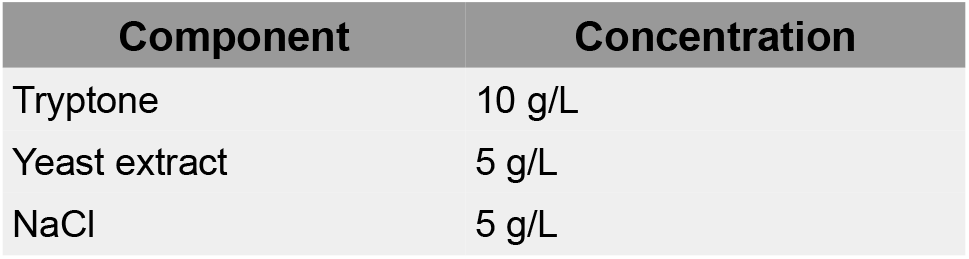

| Component | Concentration |
| --- | --- |
| Tryptone | 10 g/L |
| Yeast extract | 5 g/L |
| NaCl | 5 g/L |

All the growth media for *E. coli* are MOPS buffer based. Each media had 0.132M KH_2_PO_4_ (100X) added to it. In the case of synthetic rich media, we also added the EZ supplement and ACGU mixture. The composition of MOPS buffer, EZ supplement and ACGU mixture were followed from (*4*). Composition of each media is given below

All the *E. coli* translation inhibition experiments were done in MOPS Glucose. For the RpsB-mChartreuse reporter (SJ2524) additional translation inhibition experiments in MOPS Glycerol, MOPS Rich Glycerol and for *rpsB::mChartreuse-lacZ* (SJ2531) in MOPS Glycerol were performed.

All the *B. subtilis* media are S7_50_ salts and S7_50_ metals based (composition of these were followed from (*5*)). Each media had the following additions - 50μM iron(III) Chloride, and 50μM trisodium citrate. Also, in all the media, 40μg/mL tryptophan and 40μg/mL methionine were added for the auxotrophy of YB886 background. Composition of each media is given below

All the *B. subtilis* translation inhibition experiments were done in S7_50_ Glucose.

Antibiotic concentrations used in Fig 2, 3, and SI Fig 2.

Comparison of data generated in our study and the companion study by Li *et al* (*36*). This table is aimed to help readers navigate the two studies and provide an integrated view of the results.

### Culture procedure

Frozen glycerol stocks from -80°C were streaked on LB agar plates with corresponding antibiotics. Plates were incubated in air incubators at 37°C unless the strain contained pSIM18 plasmid in which case the plates were incubated at 30°C. Seed cultures were grown in LB lennox by suspending a colony from the plate in LB lennox. Seed cultures were grown at 37°C, shaken at 250rpm in a water bath. Seed cultures were used to inoculate pre-cultures for overnight growth in respective media at 37°C, shaken at 250rpm in a water bath. For translation-inhibition experiments, corresponding concentrations of antibiotics (chloramphenicol) were also added to pre-cultures. After 14-16 hours of growth, pre-cultures were used to inoculate prewarmed media for experimental cultures at a 100x dilution. In case of translation inhibition experiments, the antibiotics at corresponding concentration were also added in this final culture. The cultures were grown in 30mL borosilicate test tubes under the same external conditions as pre-cultures (37°C, 250 rpm). However these tubes were slanted at an angle of ∼45° to ensure proper aeration. The growth was monitored by measuring OD_600_ with a spectrophotometer at a rate of roughly two measurements per doubling. To calculate the growth rate of the cultures, a linear regime was identified in log(OD_600_) vs time plot and it was fitted with a line. The slope of this line gives the growth rate in steady-state growth of that culture.

### RNA/protein quantification

For both RNA and protein quantification, 1.5mL of culture each was collected at OD_600_∼0.4 in a centrifuge tube and instantly frozen on dry ice. OD_600_ at which samples were collected were noted down.

#### RNA quantification

The samples were defrosted in a water bath at room temperature for ∼10 minutes. The thawed samples were centrifuged and OD_600_ of the supernatant was measured -this is to track the cells lost. For the whole protocol centrifugation is done with 15000rpm for 3 minutes, unless otherwise noted. Cells were then washed twice with 600μL of ice-chilled 0.7M HClO_4_ by centrifuging and throwing away the supernatant. Then cells were digested with 300μL of 0.3M KOH for 1 hour in a 37°C water bath. The tubes were inverted every ∼15 minutes. After digestion, 100uL of 3M HClO_4_ was added to the tubes and centrifuged. The 400μL supernatant was collected in a new 2mL tube. The precipitate was then washed with 550μL of 0.5M HClO_4_ twice by centrifuging and collecting the supernatant in the 2mL tube. Then at the end we end up with 1.5mL of supernatant in the 2mL tube. This tube is centrifuged and OD_260_ of the supernatant is measured using a NanoDrop 2000 spectrophotometer. The RNA concentration (µg/ml/OD_600_) is given by OD_260_ x 31/OD_600_. The conversion factor of 31 is determined by the molar extinction coefficient of 10.5/mmol x cm and the average RNA nucleotide residue molecular weight of 324g/mol.

#### Protein quantification

The samples were defrosted in a water bath at room temperature for ∼10 minutes. The thawed samples were centrifuged at 15000 rpm for 1 minute and OD_600_ of the supernatant was measured. The centrifugation is repeated again, and after supernatant removal and measuring its OD_600_, the cells are resuspended in 200μL of water. 100μL of 3M NaOH is added to the suspension and kept at 100°C on a heat block to allow dissolution of cells and solubilization of proteins for 5 minutes. The tubes are then cooled in a room temperature water bath for 5 minutes. Then 100μL of 1.6% CuSO_4_ is added to the suspension, with vigorous shaking to mix. CuSO_4_ forms colored chelate complexes with protein peptide bonds. The tubes are left standing at room temperature for 5 minutes. After a centrifugation at 15000rpm for 3 minutes, OD_555_ of the supernatant is measured in a spectrophotometer. Alongside the cell-culture samples, 3 samples of BSA in water with 0.25mg/mL, 0.5mg/mL and 1mg/mL concentration are also processed. Plotting OD_555_ vs concentration for BSA samples gives us a conversion factor between OD_555_ and protein concentration. This factor is used to determine concentration of protein in µg/ml/OD_600_ in the cell-culture samples.

### Fluorescence microscopy and data analysis

During the steady state growth of the cultures, a 1-1.5μL sample of the culture is used for fluorescence microscopy. The 1μL sample is dropped on an agar pad made of the growth media and 2% w/v agarose (Sigma A9539-500G). In translation inhibition experiments, the agarose pad was prepared with addition of an appropriate amount of chloramphenicol. After the culture sample dries on the pad, the pad is inverted on the bottom of a wilco dish (GWST-5040, class 1.5, WilcoWells) with ∼170μm bottom thickness. Then a coverslip of 0.16 to 0.19mm thickness (VWR 48393-151) is put on the inverted pad. The sample was imaged on a Nikon Ti-E inverted microscope. For fluorescence microscopy, the source of light was Lumencore-SpectraX and the light was shone with 50% intensity. Cyan spectra from the source were used for exciting GFP and the emitted light was passed through ET-GFP (Chroma, 49002) filter. The camera used was Prime 95B sCMOS. Images were captured using 200ms exposure time. For phase contrast imaging, bright LED light was used.

In each imaging around 50-150 fields of views (FOVs) were imaged. For each FOV, a phase contrast image and a fluorescence image was captured. The images were saved as a stack as a ND2 file. Using the ND2 reader in Python, the images were separated into individual TIFF files. For cell-segmentation, Omnipose code was used (*45*). This code uses a phase contrast image and gives us a labeled image with pixels corresponding to each cell labeled. It can sometimes recognize the dirt particles on the pad or some features of the pad as cells too. So, additional filters were added to code to get rid of such non-cell objects. These filters were based on the phase contrast intensity of the midline of the object compared to the background, aspect ratio of the object, cell size below a threshold, cells lying at the boundary of the FOV and cut off due to the FOV boundaries, and finally, fluorescence intensity below a threshold. The labeled image after the filtering process was compared with the phase contrast image, manually, to make sure any object in the mask corresponding cells is not deleted or if at all, only a few cells are deleted. One such comparison is shown in Fig M1. The filtering process gets rid of almost all the dirt particles or background features that are recognized as cells.

**Fig. M1.**
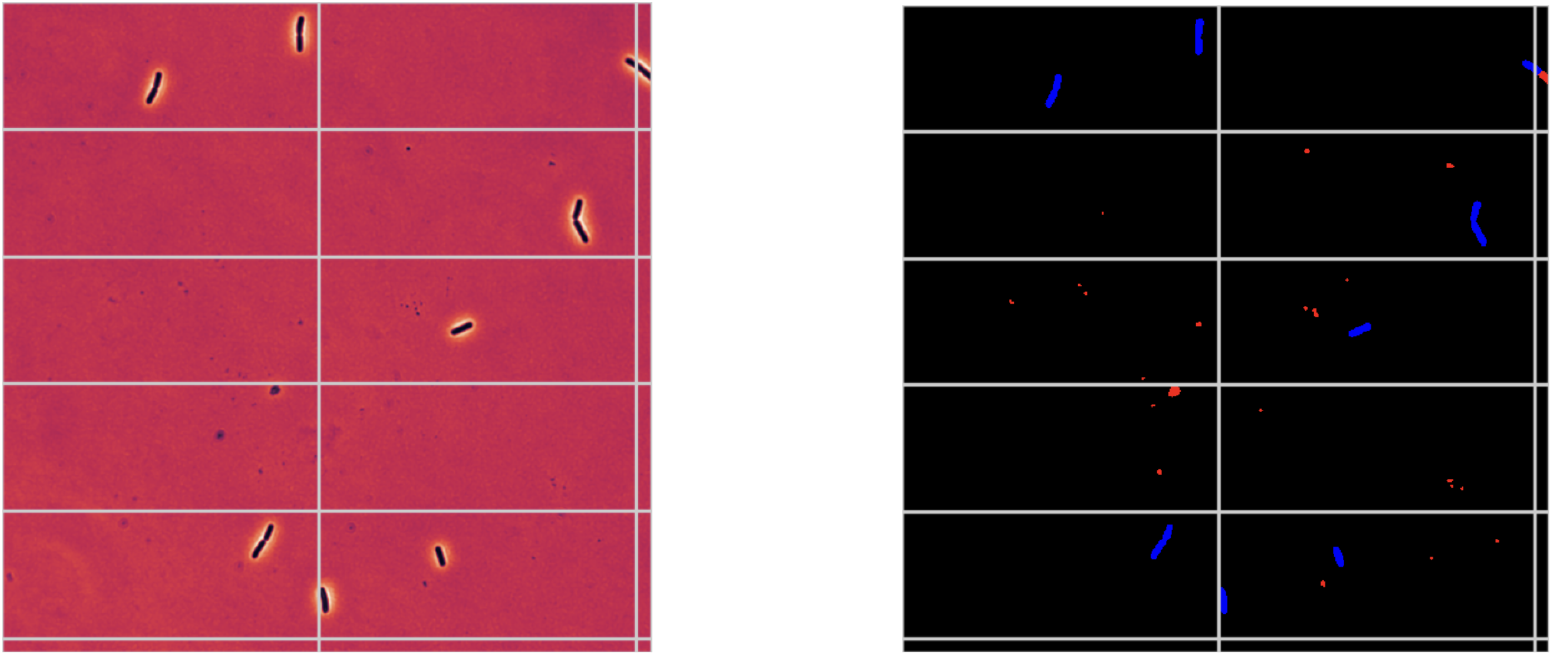
(Left) phase contrast image (Right) labeled image after filtering process. Background pixels labeled zero are shown as black color, objects that were deleted during the filtering process are colored red and objects that passed through the filter process are colored blue (used in the statistics calculation process). We can observe that the filtering process gets rid of all the dirt or background features labeled by Omnipose as cells.

With the remaining cell objects in the labeled image, various shape, size and fluorescence attributes of the cells are calculated. We calculate the average background fluorescence by summing over the fluorescence of all the pixels corresponding to the background and then dividing by the total background pixels. We subtract this average background fluorescence from each pixel corresponding to the cells. Then, we calculate the fluorescence intensity as the total background-subtracted fluorescence intensity from a cell divided by the cell volume (calculated assuming the cell as a cylinder with hemi-spherical caps). These fluorescence intensities are used in Fig. 1, 2 and 3.

### LacZ quantification and data analysis

Sample collection for LacZ quantification was done at four different OD_600_ values in steady state growth. We chose these OD_600_ values to be around 0.1, 0.2, 0.3 and 0.4. A culture aliquot of 400μL was collected per sample. The samples were collected in centrifuge tubes with 20μL of 108mM (35mg/ml) chloramphenicol already present in them. The OD_600_ at which the sample was collected was noted. The collection tubes were kept at 4°C for cooling down before sample collection. During sample collection, the culture was added to the tubes, vortexed to mix and immediately placed on dry ice. After all the samples were collected, the samples were stored at -80C for later quantification.

For quantification, the sample tubes were thawed in a room temperature water bath for ∼10 minutes. After samples were thawed, 15μL of toluene was added per tube. The tubes were vortexed to mix. Then 5μL of the sample was added to another centrifuge tube containing 495μL of Z-buffer (recipe below) + 0.27% v/v beta-mercaptoethanol. These tubes were then warmed at 37°C on a heat block for 10 minutes. Then 200μL of 2mg/mL 4-methylumbelliferyl-D-galactopyranoside (MUG, in DMSO) was added to each tube. A 250μL aliquot from each tube was added to a 96-well plate. The plate was incubated in a plate reader at 37°C for 120 minutes. A fluorescence reading, with excitation at 365 nm and emission at 450 nm, was measured every 1 minute for 3 hours, to track the product formation.

The emitted light fluorescence from a well was plotted against time to get the product formation dynamics (Fig M2-A). The initial part of this curve is linear, because substrate (MUG) is in excess compared to the LacZ molecules in the well. The linear regime was found by fitting a line to a varying set of data-points and noting the corresponding r-squared (R^2^) values (Fig M2-B). We chose R^2^ > 0.995 as the threshold for linear regime extent. Then from the **slope** of the line fit LacZ activity was calculated as below

LacZ activity (in miller units)(U/mL) = **slope** x 1000 x fold dilution in Z-buffer x 2.66

1000 × 2.66 are factors to convert the activity in miller units. Fold dilution in the Z-buffer in our case is 100. This LacZ activity is proportional to the total LacZ mass in the sample (LacZ activity = k_cat_ x total LacZ mass in the sample).

LacZ standards with known concentrations were processed in the same way as above to obtain a LacZ activity (U/mL) versus LacZ concentration (μg/mL) plot (Fig M2-C). The slope of the linear fit gives a conversion factor between LacZ activity (U/mL) and LacZ concentration (ug/mL). This factor is proportional to the catalytic activity, k_cat_, of LacZ – called as specific activity in the main text. Assuming that the catalytic activity of LacZ does not change across the growth conditions considered in the study, we use this factor to convert the LacZ activity (U/mL) in LacZ concentration (μg/mL) for all conditions.

The LacZ concentration (μg/mL) for each sample was plotted against the OD_600_ at which the sample was collected (Fig M2-D). This gives a linear plot passing through the origin, the slope of which is LacZ concentration per biomass for the given growth condition (μg/mL/OD). This was then divided by the protein concentration (μg/mL/OD) obtained in the same growth condition using the biuret method as explained above, which then gives us LacZ mass as a fraction of total protein mass [LacZ] (μg/μg) which is plotted in Fig 3.

**Fig. M2.**
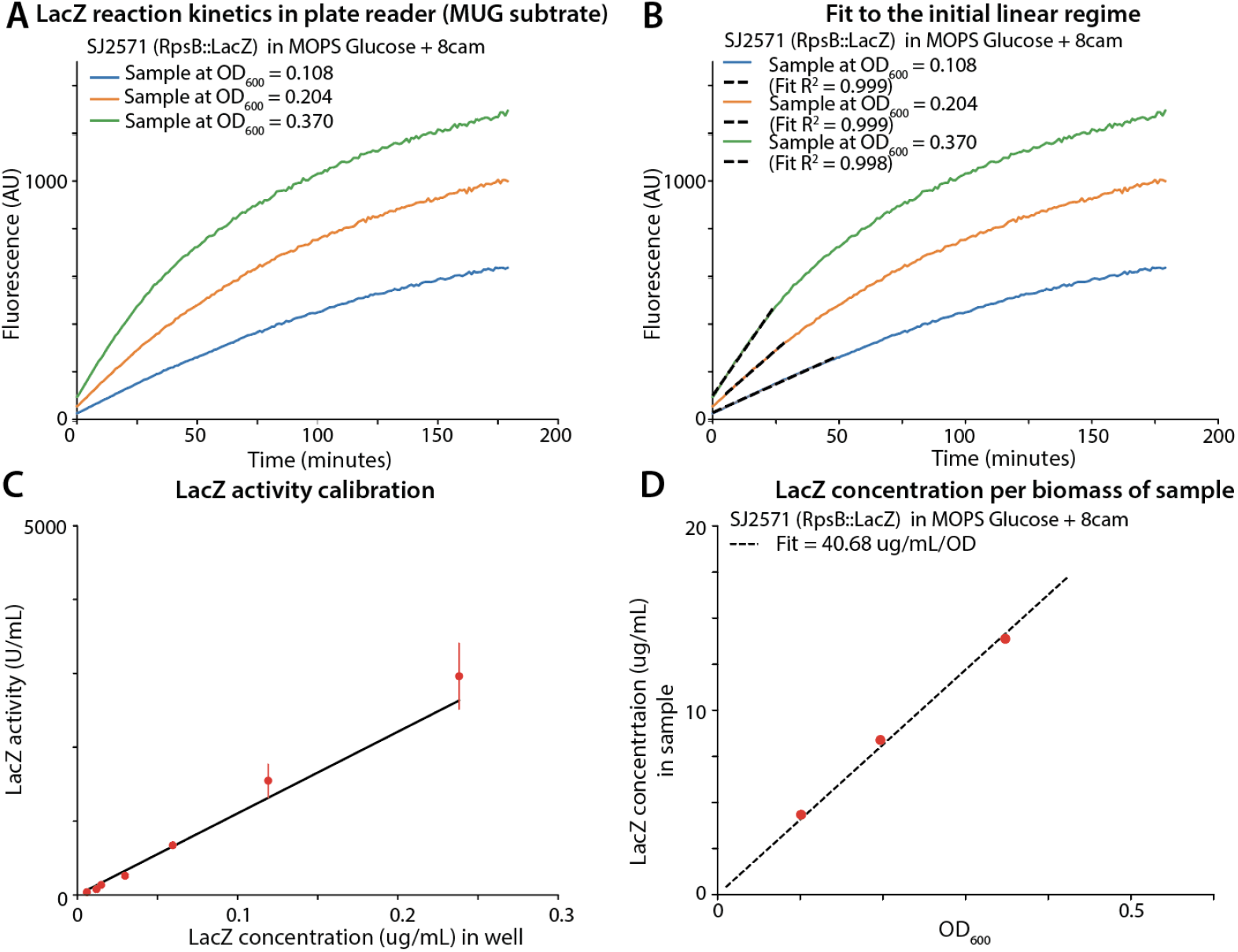
(A) LacZ reaction in plate reader; Fluorescence measured at excitation wavelength 365nm and emission wavelength 450nm, (B) Linear fits to the linear regime at the beginning of the reaction when MUG substrate is in significantly high amount. Cutoff of the linear regime was when R^2^ of the fit is >0.995. The slope of the fit is used to calculate LacZ activity (U/mL) (C) LacZ activity (U/mL) plotted versus concentration of the lacZ standards in the well plate. The slope of this line gives a conversion factor between LacZ activity (U/mL) and LacZ concentration (μg/mL). This conversion factor is used to convert activity calculated in panel B to LacZ concentration in the sample (μg/mL) (D) LacZ concentration in the sample plotted against the OD_600_ at which the sample was collected. The slope of this line gives LacZ concentration per biomass (μg/mL/OD).

## Acknowledgements

This research was supported by the National Institutes of Health (grant award no R35GM139622 to S.J.), a grant from the Simons Foundation (SFI-PD-Pivot Fellow-00008375 to S.J.), and a grant from the Agence Nationale de Recherche (PEPR B-BEST, ANR-24-PEBB-0006 to H.d.J). We also thank Danny Fung (Jue Wang Lab, UW Madison) for construction of *B. subtilis* RpsB-mChartreuse reporter strain.

## Data and Material Availability

All the data are present in the materials and methods, and supplementary materials. All the materials can be provided on a reasonable request to Suckjoon Jun

## Supplementary Materials

SI Note 1

Supplementary Figures 1 and 2

Data S1

## Supplementary Material

**SI note 1 - Fluorescence intensity and LacZ activity from the reporters are expected to be proportional to the underlying ribosomal abundance (RNA-to-protein mass ratio)**

**Definitions**

In the introduction section, we defined ribosomal abundance as the total ribosomal protein mass (M_rib_) as a fraction of total protein mass (M_protein_) in a cell:

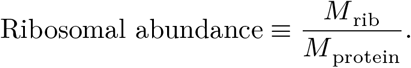

We now define the RNA-to-protein ratio (ρ) and fluorescence intensity (f) of a cell as follows :

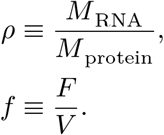

Here, M_RNA_ and M_protein_ denote total cellular RNA and protein mass, respectively. F is the background-subtracted total fluorescence intensity of a cell and V is the volume of the cell.

Following are some quantifies used in the derivations below:

| Symbol | Meaning |
| --- | --- |
| $m_{\text{rib}}$ | Mass of ribosomal proteins in a single *E. coli* ribosome |
| $m_{\text{rRNA}}$ | Mass of rRNA per ribosome |
| $m_{\text{mChartreuse}}$ | Mass of a single mChartreuse molecule |
| $N_{\text{RspB}}$ | Total number of RpsB molecules in an *E. coli* cell during steady state growth |
| $N_{\text{rib}}$ | Total number of ribosomes in an *E. coli* cell during steady state growth |
| $N_{\text{mChartreuse}}$ | Total number of mChartreuse molecules in an *E. coli* during steady state growth |
| $M_{\text{rib}}$ | Mass of all ribosomal proteins in an *E. coli* cell |
| $M_{\text{RNA}}$ | Mass of all RNAs in an *E. coli* cell |
| $M_{\text{rRNA}}$ | Mass of all rRNAs in an *E. coli* cell |
| $M_{\text{protein}}$ | Mass of all proteins in an *E. coli* cell |
| $M_{\text{cell}}$ | Dry mass of an *E. coli* cell |
| $R_{\text{tr}}$ | Transcription rate from a promoter in mass of mRNA generated per unit time |
| $R_{\text{tl}}$ | Translation rate of a transcript in mass of peptide chain generated per unit time |
| $R_{\text{d}}$ | Degradation rate of an mRNA/rRNA transcript |
| $[g]$ | Gene number concentration in the cell |
| $\epsilon$ | Fluorescence output of a single mChartreuse molecule (even when in fusion with LacZ in the dual reporter, mChartreuse-LacZ) |
| $C$ | DNA replication period in *E. coli* |
| $V$ | Volume of an *E. coli* cell |
| $\lambda$ | Growth rate of an *E. coli* cell |

### Relation between ribosome abundance and RNA-to-protein ratio (ρ)

Around 86% of RNA in exponentially growing *E. coli* cells is ribosomal RNA (rRNA) across multiple growth conditions (7):

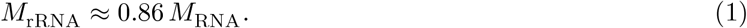

In addition, ribosomal proteins and rRNA are produced in stoichiometric proportion (7, 8, 17):

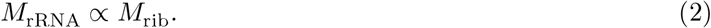

Combining equations (1) and (2) gives

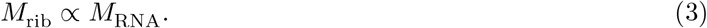

In exponentially growing cells, cell density is approximately constant across multiple growth conditions (46), implying

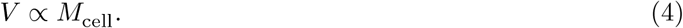

The protein fraction of biomass is also approximately constant over a range of growth conditions (38), so

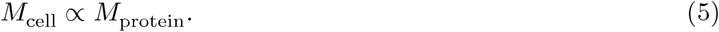

Combining equations (4) and (5) gives

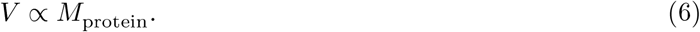

Equation (6) therefore implies

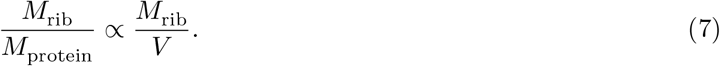

Thus, ribosomal abundance 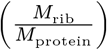 is proportional to the ribosomal-protein concentration 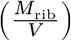 in an *E. coli* cell.

Using equation (3), we also obtain

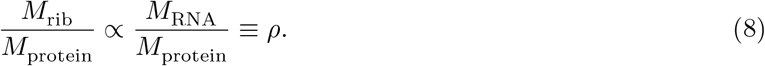

Hence, ribosomal abundance is proportional to the RNA-to-protein ratio ρ. As all the relations above hold across range of growth conditions, this relationship holds true for all the growth conditions studied here.

### Relation between RpsB-mChartreuse fluorescence intensity (*f*) and RNA-to-protein ratio (ρ)

We assume that each mChartreuse (GFP) reporter contributes fluorescence ϵ. We call this the fluorescence output per reporter molecule in the main text. We also assume that the RpsB–mChartreuse fusion does not affect ribosome assembly, consistent with growth-phenotype data from earlier studies using RpsB–GFP fusions (28, 31). Because each ribosome contains one RpsB molecule, each ribosome is associated with one mChartreuse molecule.

The total cellular fluorescence is therefore

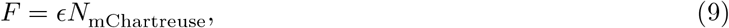

where *N*_mChartreuse_ is the number of mChartreuse molecules in the cell. We also have

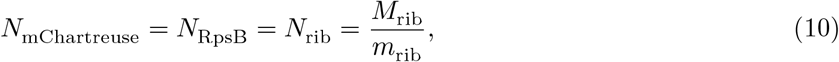

where we have used a single mChartreuse molecule per ribosome assumption.

Combining (9) and (10) with equation (7) gives

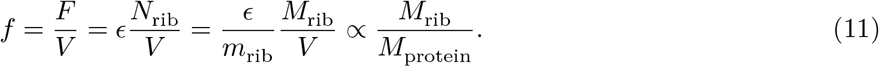

That is, the fluorescence intensity is expected to be proportional to the ribosomal abundance, under the assumption that the fluorescence output per unit reporter is constant. This along with equation (8) gives

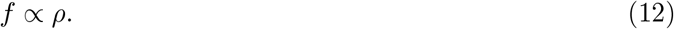

Hence, fluorescence intensity from the RpsB-mChartreuse translational fusion is expected to be directly proportional to the RNA-to-protein ratio.

### Relation between fluorescence intensity (*f*) and RNA-to-protein ratio (ρ) in *rpsB::mChartreuse* and *rpsB::mChartreuse-lacZ* ; also between [LacZ] and RNA-to-protein ratio (ρ) in *rpsB::mChartreuse-lacZ*

We assume again that each mChartreuse molecule contributes fluorescence ϵ.

In strains *rpsB::mChartreuse, rpsB::lacZ*, and *rpsB::mChartreuse-lacZ, mChartreuse, lacZ*, and *mChartreuse-lacZ* reporter constructs are downstream of the *rpsB* gene. However, both the reporter construct and the *rpsB* gene share the same promoter. Hence, both will have the same rate of transcription (*R*_tr_). By construction mChartreuse in SJ2570 (*rpsB::mChartreuse*), LacZ in SJ2571 (*rpsB::lacZ*) and mChartreuse-LacZ in SJ2574 (*rpsB::mChartreuse-lacZ*) have the same ribosome binding site and hence the same translation rate (*R*_tl_). RpsB, however, will have a different translation rate because it has a different ribosome binding site.

The steady-state concentration of RpsB, mChartreuse, LacZ and mChartreuse-LacZ protein molecules would be given by solving a system of coupled ODEs at steady state, describing the rate of change of mRNA concentration:

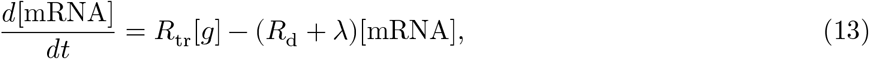

and rate of change of protein concentration

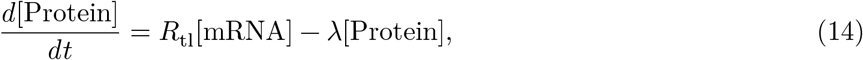

where parameters are defined in the table above. These equations apply for RpsB, mChartreuse, LacZ and mChartreuse-LacZ reporter molecules, which share the same *R*_tr_, *R*_d_, but have different *R*_tl_. For mRNA, the degradation rate (*R*_d_) is much higher than the dilution rate due to the growth (λ), so that we can ignore the growth dilution of mRNA. We have also ignored the protein degradation rate, for it is negligible with respect to growth dilution in the conditions considered here.

Then solution at steady state gives the RpsB concentration ([RpsB]) as

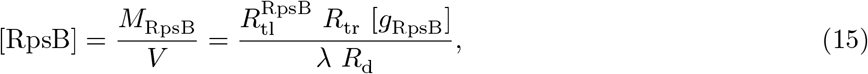

where 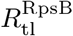 is the translation rate for RpsB and [*g*_RpsB_] is the gene concentration of RpsB. The concentration of mChartreuse ([mChartreuse]) in SJ2570 (*rpsB::mChartreuse*) is given by

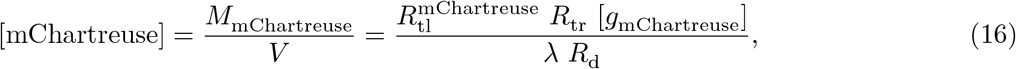

The solution should be analogous for [LacZ] and [mChartreuse-LacZ] :

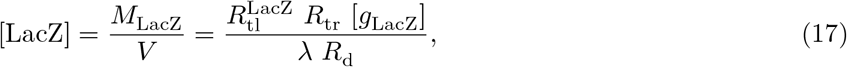

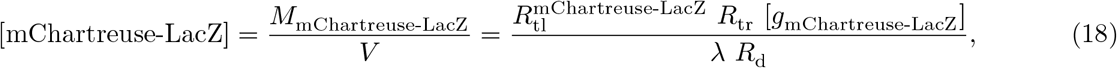

Because mChartreuse and RpsB are in the same operon in SJ2570,

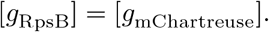

Similarly for LacZ in SJ2571 and mChatreuse-LacZ SJ2574, we have

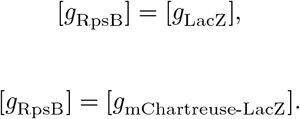

Even though mChartreuse and RpsB have different ribosome binding sites in SJ2570, we can assume that their translation rates are proportional to each other across growth conditions, so that

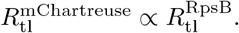

Similarly for SJ2571 and SJ2574,

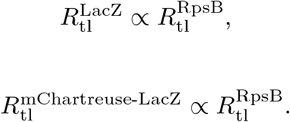

Then equations (15), (16), (17) and (18) imply

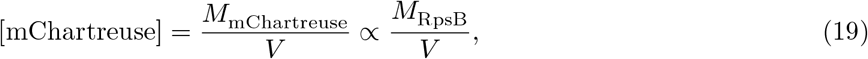

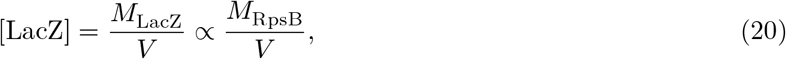

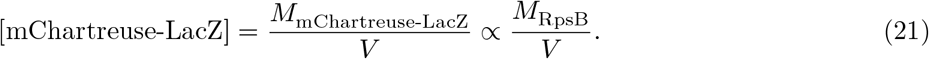

Then for the fluorescence intensity (*f*) from mChartreuse in SJ2570, we get

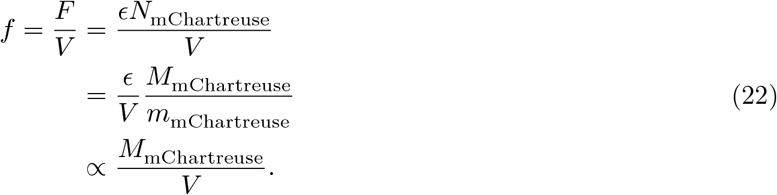

A similar argument should follow for mChartreuse-LacZ in SJ2574.

Because there is a single RpsB molecule per ribosome,

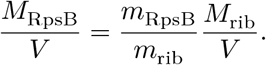

This, along with equations (19), (20), (21), and (22), implies

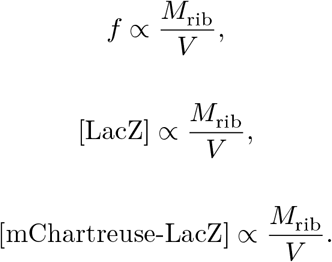

Hence fluorescence intensity and LacZ concentrations are proportional to ribosome concentration, which along with equations (7) and (8), give

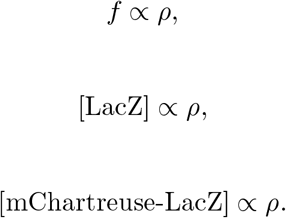

Note here, we defined [LacZ] as mass concentration - but using equation (6) we get that LacZ mass concentration is proportional to LacZ mass as a fraction of total protein mass, as defined in the main text.

### Relation between mChartreuse fluorescence intensity (*f*), [LacZ] and RNA-to-protein ratio (ρ) in ectopically expressed PrrnB::*mChartreuse*, PrrnE::*mChartreuse*, and PrrnB::*mChartreuse-lacZ* strains

Assume that each mature mChartreuse molecule contributes fluorescence ϵ.

The native *rrnB* operon and the ectopic *mChartreuse* reporter gene are controlled by the same *rrnB* promoter sequence. Therefore, their transcription rates per gene copy are assumed to be proportional:

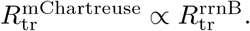

For the PrrnE::*mChartreuse* construct, the corresponding assumption is

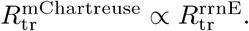

Following equation (13), the steady-state cellular concentration of rRNA transcribed from the native *rrnB* operon is

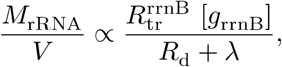

where 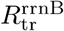 is the transcription rate per native *rrnB* gene copy, [*g*_rrnB_] is the gene concentration of the native *rrnB* operon, *R*_d_ is the rRNA degradation rate and λ is the growth rate. The proportionality sign, instead of equality, is because rRNA expressed from other *rrn* operons also contribute to *M*_*rRNA*_. Because rRNA is stable and only a small fraction of it is degraded in growth conditions of this study, *R*_d_ is assumed negligible with respect to λ. Then we get,

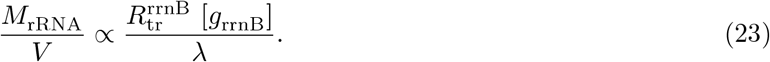

Again following equations (13), (14) and short-half life of mRNA, the steady-state mChartreuse protein concentration is

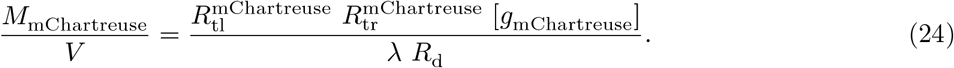

The relationship between the native *rrnB* gene concentration and the ectopic reporter-gene concentration is written as (47)

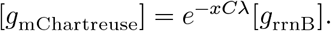

Similarly for rrnE reporter strain

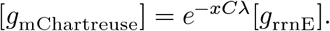

where *C* is the period DNA replication in *E. coli*, and *x* is the coordinate of the ectopic site, defined such that *x* = 0 is *oriC* and *x* = 1 is *ter*.

Transcriptional activity of the ectopic PrrnB reporter is expected to be approximately the same as transcription from the native rrnB operon, so that

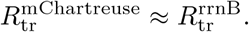

With the above relations and the assumption that the translation rate 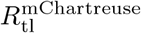 and degradation rate *R*_d_ do not change across the growth conditions, equation (24) gives

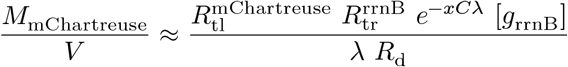

Then using equation (23) we have

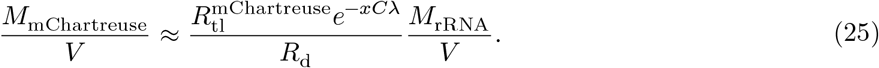

In our case, *x* = 0.64 for SJ2560 (PrrnB::*mChartreuse*) and SJ2561 (PrrnE::*mChartreuse*) strains, which have the insertion near the *tolA* gene and *x* = 0.75 for SJ2574 (PrrnB::*mChartreuse-lacZ*) strain, which has the insertion is near the *appA* gene. We assume that *C* = 40 minutes = 0.667 hrs (4, 48) across all the nutrient-limited growth conditions. Thus, between the lowest (λ = 0.25/hr) and highest (λ = 1.25/hr) nutrient-limited growth rates in our study, the exponential factor *e*^−*xC*λ^ leads to a 35-40% change. This change is smaller compared to the more than 400% change observed in the activity of PrrnB and PrrnE promoters (20, 32). During chloramphenicol inhibition, Si *et al*. (4) report a 4 times increase in *C* between 0 μM to 10 μM, however concomitantly there is a 4-fold decrease in growth rate (λ). This makes the exponential factor *e*^−*xC*λ^ approximately constant over the chloramphenicol treatment range. Following these two arguments we can absorb the exponential factor into the proportionality. Then we get from equation (25):

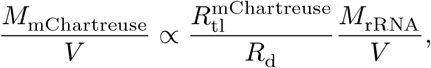

and using equations (1) and (6),

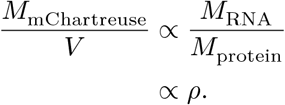

Following the arguments of previous sections,

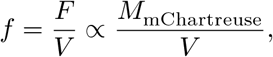

and hence,

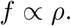

This argument similarly applies to fluorescence intensity (*f*) and [mChartreuse-LacZ] in SJ2574 (PrrnB::*mChartreuse-lacZ*).

## Supplementary figure 1

**Figure S1.**
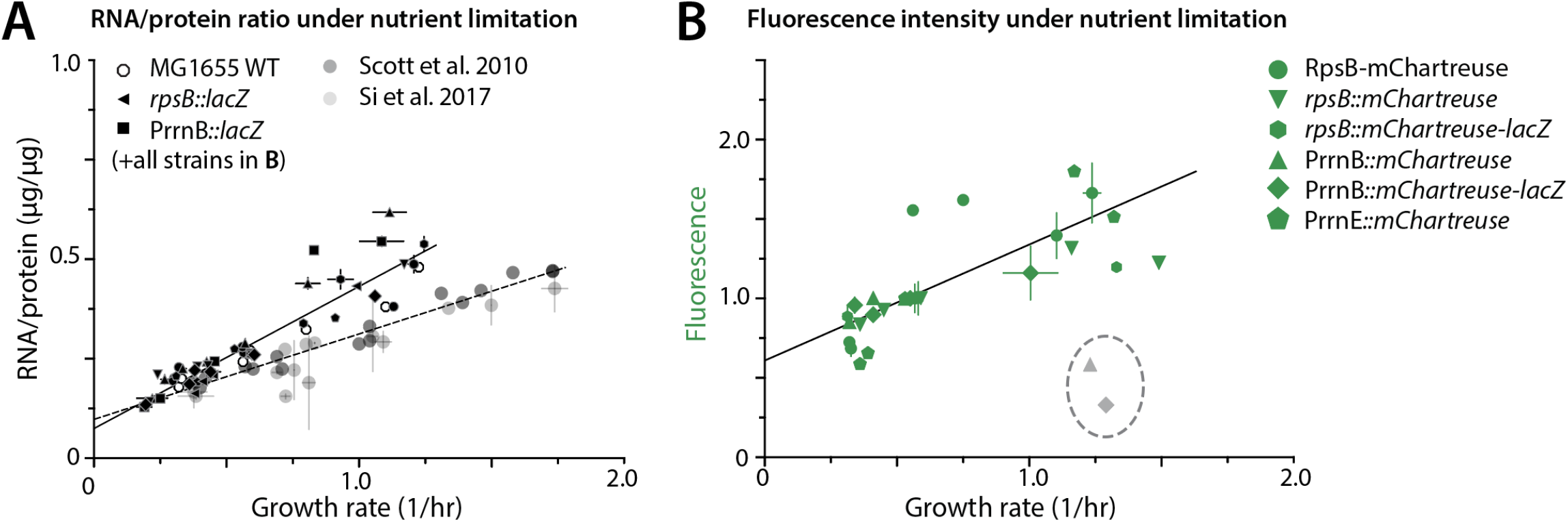
RNA-to-protein mass ratio and Fluorescence intensity increase linearly with growth rate under nutrient limitation. (**A**) RNA-to-protein mass ratio (RNA/protein) against growth rate under nutrient-limited growth, for wild-type MG1655, the *lacZ* reporter strains, and all mChartreuse reporter strains listed in **B.** Grey symbols are previously published measurements (3, 4). Solid line, fit to the strains of this study; dashed line, fit to the published data. (**B**) Fluorescence intensity against growth rate under the same conditions. Fluorescence is normalized to the value measured in MOPS glucose for each strain and hence it is unit less. Grey points are fluorescence data for PrrnB::*mChartreuse* and PrrnB::*mChartreuse-lacZ* in MOPS Rich Glycerol. For these growth conditions, the fluorescence values were much lower than other data points for the same strains. We are unable to explain this unexpected drop in fluorescence in MOPS Rich Glycerol. They were not included in the fit. Error bars are calculated as standard error of the mean between at least two replicates. Replicates information is given in Data S1.

## Supplementary figure 2

**Figure S2.**
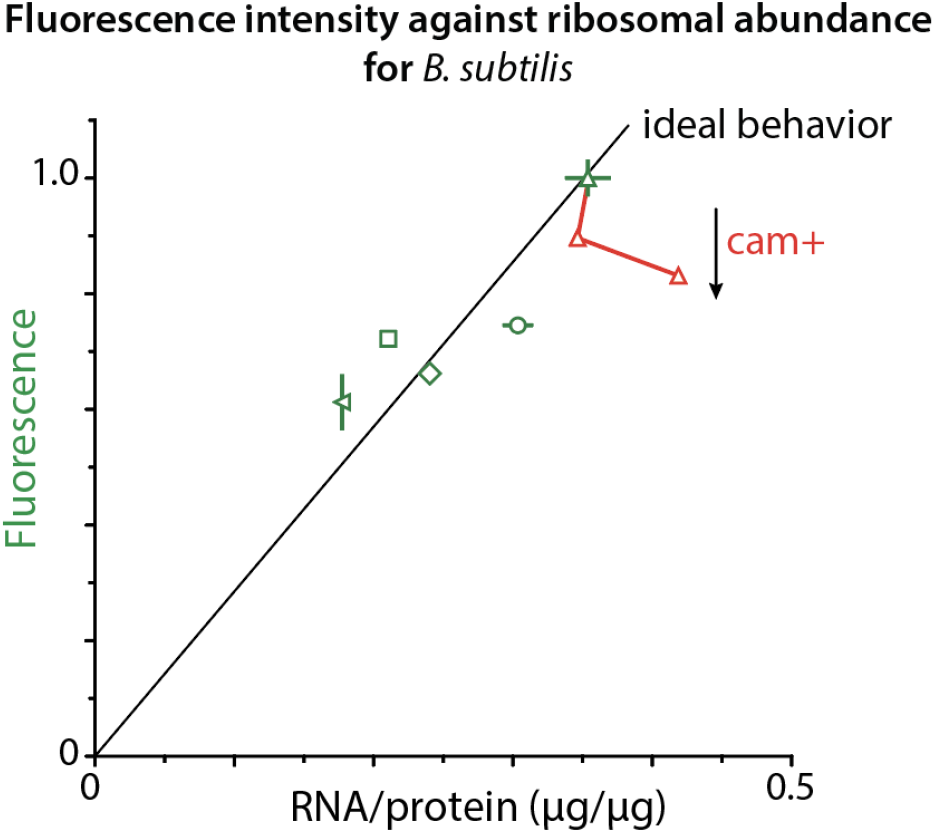
Fluorescence intensity and RNA/protein relation for *B. subtilis* RpsB-mChartreuse. The growth media are given in Table 3. The ideal behaviour line is fit to the data forcing the intercept zero. Chloramphenicol was added to S7_50_ Glucose with concentrations 4μM and 6μM. Error bars are calculated as standard error of the mean between at least two replicates. Replicates information is given in Data S1.

## Data S1

Data generated in this study. For all the strains, data across different growth media and chloramphenicol (cam) concentrations are provided. In each growth condition, Average (Avg), Standard error of mean (SEM), and number of replicates (Replicates) are provided for Growth rates (1/hr), RNA/protein (μg/μg) values, Fluorescence intensities (normalized by MOPS Glucose fluorescence intensity), and [LacZ] (μg/μg) values.

## Notes

### Competing Interest Statement

The authors have declared no competing interest.

### Summary of Updates

Added supplementary data set, added reference to a companion study which has been submitted as a preprint on bioRxiv (https://doi.org/10.64898/2026.09.30.755718), updated paragraph 2 on page 3, added data and material availability statement and supplementary material information at the end of the main text, all the error bars in the figure are made standard error of mean

